# Rhodamine-derived ratiometric fluorescent molecular rotor for mitochondrial viscosity sensing

**DOI:** 10.64898/2026.09.13.751207

**Authors:** Oksana Kharchenko, Pascal Didier, Andrey S. Klymchenko

## Abstract

Fluorescent molecular rotors for sensing local microviscosity have become indispensable tools for better understanding the mechanisms of living systems. Current probes relay on intensity or lifetime measurements, while ratiometric molecular rotors are lacking. Here, we introduce a new design concept of molecular rotor using xanthene scaffold with freely rotating 9-aryl group. We studied three rhodamine derivatives, where 2-carboxyphenyl group was replaced with 4-methoxyphenyl, 2-thienyl and 2-benzofuryl. We found that five membered 9-aryl ring provides two major effects. First, free rotation of thienyl derivative, confirmed by theoretical calculations, makes it classical molecular rotor with intensity and lifetime-based response to viscosity. Second, benzofuryl derivative exhibits second emission band in near-infrared region and fluorescence ratiometric response to viscosity. Theoretical calculations suggest that the benzofuryl derivative can attain planar conformation in the excited state, which explains appearance of the NIR emitting band. Both new dyes efficiently target mitochondria, while the 9-benzofuryl derivative enable quantitative ratiometric measurement of viscosity of inner mitochondrial membrane. Ratiometric imaging revealed mitochondria heterogeneity with local viscosity values ranging from 250 to 460 cP. Oxidative stress induced significant rise in the local viscosity in mitochondria in addition to their morphological changes. Overall, we propose a concept that converts bright rhodamine dyes into molecules rotors with a valuable ratiometric response to viscosity, which opens a bunch of potential biological applications.

## Introduction

Sensing molecular environment has been established as a powerful concept for imaging and sensing. It is realized by so-called environment-sensitive fluorescent probes capable of changing their fluorescence intensity and color in response to their environment polarity, hydration, viscosity, electric fields, etc.^1-4^ On the one hand, those probes can provide the information about the molecular organization of biomolecules, such as lipids in biological membranes, proteins in their aggregates or nucleosides inside DNA/RNA. On the other hand, they are universal tools that sense practically any type of biomolecular interactions, including protein-protein, protein-membrane, protein-nucleic acid, ligand-receptor, etc. Therefore, there is an unmet need for the development of new advanced environment-sensitive probes featuring higher brightness, photostability, sensitivity to the environment and operation in favorable red region of visible spectrum.

The environment-sensitive probes could be classified into solvatochromic dyes,^1,5^ molecular rotors^1,6^ as well as dyes undergoing ground-state conformational change (e.g. flippers and butterflies)^7^ and isomerization (e.g. spironolactone-forming rhodamine derivatives).^8-10^ Among these dyes, molecular rotors are the tools of choice when it comes to sensing local viscosity in biological structures or materials.^1,6^ The early examples of molecular rotors were introduced by Haidekker and coworkers based on push-pull dye 9-(dicyanovinyl) julolidine (DCVJ) and its ester analogues.^6,11^ A large variety of molecular rotors developed to date rely on the similar push-pull design principle, which ensures twisted intramolecular charge transfer (TICT),^12^ the process that strongly depends on the viscosity. The prominent examples include ThT, extensively used to study protein aggregation,^13^ GFP fluorophores that light up in highly rigid environments of RNA aptamers^14^ and engineered proteins.^15^ Red-shifted absorption and improved brightness in this type of dyes can be exemplified by dioxaborine-based dye DXB-Red, which was applied for sensing ligand-receptor binding.^16^ However, this design concept has a fundamental limitation – the dyes present sensitivity to both polarity and viscosity. Moreover, TICT based quenching is most efficient in polar media, which means that these probes are more efficient viscosity sensors in polar environment.^16^ The breakthrough in the field was made with introduction of BODIPY molecular rotor by Kuimova and co-workers.^17^ In this case, the design is based on conjugation of bright fluorophore with an aromatic group that undergoes free rotation leading to appearance of pathways for non-radiative deactivation. BODIPY rotors unlike TICT-based probes exhibit clean response to viscosity with nearly no response to solvent polarity. On the other hand, it inherits favorable properties of BODIPY in terms of brightness and photostability. Therefore, BODIPY rotor found impressive number of applications in multiple biological fields,^4^ which includes sensing viscosity and lipid order in biological membranes^18-20^ and monitoring protein aggregation in vitro^21-22^ and tumor microscopic viscosity in vivo,^23^ and microviscosity mapping in plant cells and tissues,^24^ and temperature sensing in organelles. Thus, its derivatives bearing targeting groups were successfully applied to study local viscosity and lipid organization or protein nano-environment of different organelles, including mitochondria.^25-28^ However, this dye presents several major limitations. First, extension of BODIPY rotor emission to red region made them sensitive to temperature rather than viscosity.^29^ Second, this probe operates by changes in the fluorescence intensity, which is difficult to quantify using fluorescence imaging because the intensity depends on local probe concentration and multiple instrumental factors. Therefore, it is mainly used for sensing viscosity by measuring changes in its fluorescence lifetime, in particular using fluorescence lifetime imaging (FLIM).^4^ Examples of ratiometric probes to viscosity are quite rare and limited to dye dyads. Thus, FRET-based ratiometric molecular rotors were developed where the FRET donor operates as a reference (viscosity insensitive unit emitting in blue), while FRET acceptor (emitting in red) is sensitive to viscosity.^30^ The other example is dyad of two porphyrins connected by a conjugated linker. It presents several emissive conformation states, sensitive to local viscosity, leading to viscosity-dependent dual emission.^31^ However, these types of probes are relatively large molecules that are not always well suited for biological applications, which generally require compact probes. Moreover, dyads can exhibit complex photophysical behavior and independent photobleaching of the two chromophores, which may lead to drastic changes in their spectral properties and, consequently, to measurement artifacts, especially in fluorescence microscopy, where relatively high excitation powers are often used. In this respect, we turned our attention to xanthene derivatives, which gave rise to commonly used fluorescein and rhodamine dyes. Early works showed that removal of carboxylate group from fluorescein decreased the fluorescence quantum yield (QY),^32^ which was later explained by the non-radiative deactivation related to the rotation of phenyl group in 9-position.^33^ Similar phenomenon was observed when 2’-methyl group was removed from Tokyo Green,^34^ confirming the importance of free rotation of 9-phenyl group in quenching fluorescence of fluorescein dyes. Decrease in the rotation barrier is an essential factor impacting sensitivity of dyes to viscosity.^35^ However, another study dedicated to rhodamine and rosamine dyes showed that removal of 2’-subsistutent does not produce any effect of emission of rhodamine analogues.^36^ On the other hand, another study showed that the presence 4’-dialkyl amino group in the rosamine dye leads to strong fluorescence quenching and “bi-chromophore effect” with appearance of new long-wavelength emission bands, where an excited state involving 9-phenyl group is considered.^37^ Further study an additional phenyl ring was introduced, which led to a fluorogenic dye that increased its emission in more rigid environments of macrocycle.^36^ A very recent dye by Kikuchi and co-workers proposed to decrease the steric hindrance in the 2-aryl group by replacing it with furyl and thienyl groups.^38^ The obtained dyes showed typical molecular rotor properties with large variation of the fluorescence quantum yield as a function of viscosity. However, this system behaved like BODIPY rotor, with a intensity and lifetime response without the ratiometric output.

Based on these reports, we hypothesized that 9-aryl group containing electron rich 5-membered ring with extended conjugation could combine free rotation with respect to xanthene fluorophore and possibility of creating “bi-chromophore effect”.

In the present work, we synthesized a series of rhodamine derivatives bearing five membered rings: thienyl and benzofuryl as well as reference dye with 4’-methoxyphenyl group in the 9-position. We found that both dyes with 5-membered rings exhibit much stronger fluorescence quenching in low-viscosity solvents and they exhibited a significant enhancement of their emission intensity and the lifetime with viscosity, similar to that observed for BODIPY rotor. Importantly, benzofuryl derivative of rhodamine displayed in low-viscosity solvent a second emission band in the far-red region, where the emission ratio of these two emission bands varied with solvent viscosity. This is one of the first examples of ratiometric viscosity probe that operates based on a single fluorophore. Cellular studies revealed that it targets mitochondria and reports on the local changes in the mitochondrial membrane organization in response to multiple stress conditions, such as oxidative stress. The developed dyes constitute unique tools for monitoring local viscosity in biological systems and materials science using simple ratiometric measurements in fluorescence spectroscopy and microscopy.

## Results and discussion

### Design and synthesis

Three rhodamine derivatives were synthesized: Ph-R, Th-R and Bf-R (Figure 1). As the aryl group at the 9-position of xanthene is expected to play a key role in the control of the fluorescence quantum yield, we made two types of changes with respect to parent rhodamine B. First, we replaced 2-carboxyphenyl group with 4-methoxyphenyl to obtain a rosamine derivative Ph-R. Second, we replaced phenyl group in rosamine dye with thienyl and benzofuryl groups. We expected that the steric hindrance for the rotation would decrease in the following sequence: Rh-R > Th-R > Bf-R (Figure 1). Indeed, five-membered ring of benzofuryl heterocycle should provide minimal sterical hindrance because its oxygen atom is smaller compared to sulfur of the thienyl group. Moreover, in case of in benzofuryl heterocycle, fused benzene ring could provide addition stabilization of the planar form of the dye with longer electronic conjugation. Previous works suggested that electron donor 9-aryl group (dialylaminophenyl) can participate in the excitation, providing addition emission band at longer wavelength in low temperature glasses.^37^ Our rosamine dye Ph-R was designed to bear para-methoxy group, to ensure a weak electron donor properties of aryl group similar to those in Th-R and Bf-R. In this study, it serves as reference molecule for Th-R and Bf-R, in order to understand the effect of five membered ring and the electronic conjugation on the xanthene scaffold. All three compounds were synthesized in two steps starting from the reaction of 3-(diethylamino)phenol with corresponding aromatic aldehyde, followed by oxidation step with chloranil. They were purified by column chromatography identified by NMR and mass spectrometry.

**Figure 1.**
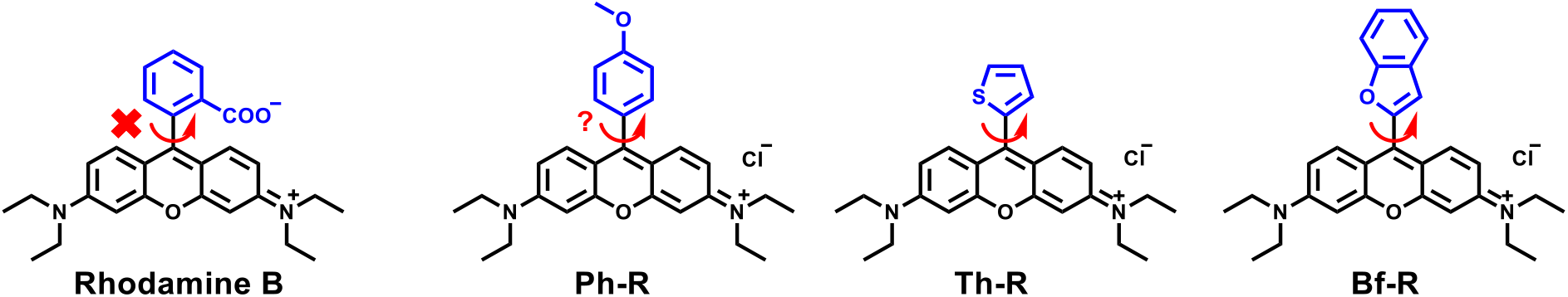
Rhodamine B and the newly synthesized derivatives of the presented study. Arrow shows impossible rotation of 9-aryl group in rhodamine B, questionable rotation in Ph-R and potential rotation for five-membered 9-aryl groups of Th-R and Bf-R.

### Spectroscopic properties in organic solvents

We first characterized their absorption and fluorescence properties in organic solvents (Figure 2). The absorption band of the dyes showed systematic red shift in the order Rh-R → Th-R → Bf-R, which overall shift from 553 nm for the reference dye Ph-R to 590 nm for Bf-R in methanol. Given that the aryl groups in all three dyes present comparable weak electron donor ability, this systematic red shift suggests an increase in the electronic conjugation between 9-aryl and xanthene moiety. The polarity of the solvent did not show significant effect on the position of the absorption maximum. Emission spectra of thienyl also displayed red shift compared to Ph-R, which corresponded to that observed in the absorption spectra (Figure 2). Surprisingly, the emission spectrum of Bf-R was completely different because it contained a highly emissive red shifted band at 725 nm as well as short-wavelength shoulder (Figure 2), which probably corresponded to “classical” emission band of a rosamine derivative. To verify whether both bands originate from the same ground state, we recorded the excitation spectra at both emission bands. The obtained two excitation spectra were nearly the same and matched closely the absorption spectrum of the dye, indicating that the two emission bands originate from some process in the excited state. The fluorescence quantum yields (QY) of the Ph-R were high (50-94%) for most of the solvents (Table 1). In contrast, the QY of and Th-R and Bf-R dyes were systematically lower than those of Ph-R in all studied solvents. Thus, five-membered heterocycles provoke fluorescence quenching of the new dyes.

**Table 1.** Spectroscopic properties of Ph-R, Th-R and Bf-R dyes.^a^.

| Dye | Parameters | Solvents or media |  |  |  |  |  |
| --- | --- | --- | --- | --- | --- | --- | --- |
|  |  | PB | DMSO | ACN | MeOH | Acetone | THF |
| **Ph-R** | $\lambda_{\text{Abs}}$ | 555 | 563 | 553 | 553 | 555 | 557 |
| | $\lambda_{\text{Em}}$ | 579 | 590 | 578 | 575 | 579 | 579 |
|  | QY, % | 50 | 94 | 58 | 86 | 61 | 94 |
| **Th-R** | $\lambda_{\text{Abs}}$ | 574 | 583 | 572 | 571 | 574 | 576 |
| | $\lambda_{\text{Em}}$ | 598 | 613 | 600 | 597 | 602 | 602 |
|  | QY, % | 5 | 35 | 12 | 24 | 14 | 17 |
| **Bf-R** | $\lambda_{\text{Abs}}$ | 608 | 603 | 590 | 590 | 592 | 595 |
| | $\lambda_{\text{Em}}$ | - | 745 | 723 | 723 | 723 | 723 |
|  | QY, % | - | 23 | 25 | 15 | 21 | 24 |

<sup>a</sup> $\lambda_{\text{Abs}}$ and $\lambda_{\text{Em}}$ are absorption and emission band maxima. QY is a relative fluorescence quantum yield.

**Figure 2.**
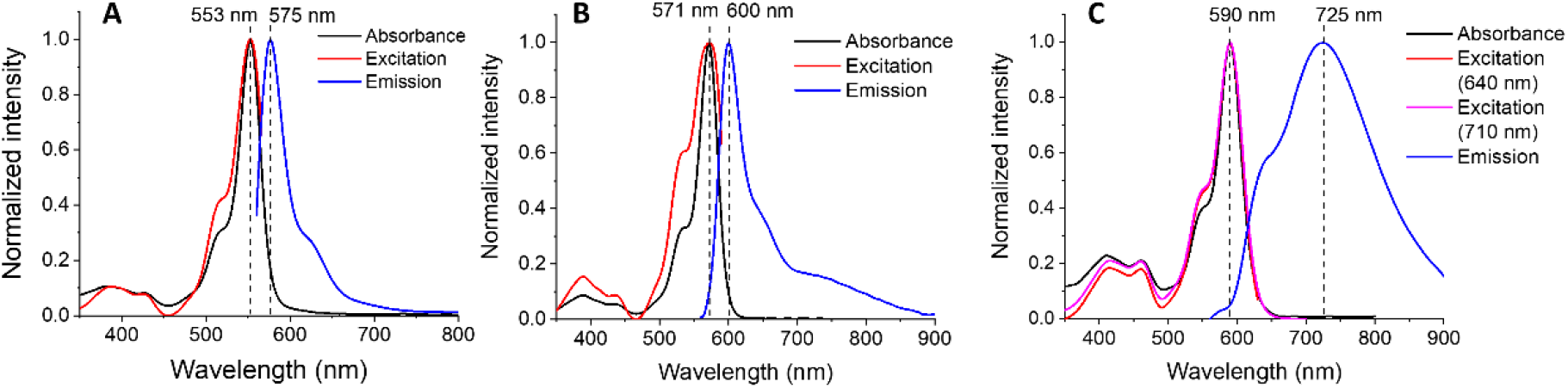
Normalized absorbance, excitation, and emission spectra of (A) Ph-R, (B) Th-R and (C) Bf-R recorded in MeOH. λ_ex_ = 550 nm.

Next, we studied the effect of viscosity for the three dyes in methanol-glycerol mixtures. The reference compound showed only minor increase in the fluorescence intensity upon increase in the glycerol content, indicating poor molecular rotor properties (Figure 3), in line with previous reports.^38^ In sharp contrast, Th-R showed strong increase in the fluorescence intensity upon increase in viscosity. As a result, the quantum yield in pure glycerol was close to 100%. Thus, replacing six-membered phenyl group with five-membered thienyl group transforms the dye into molecular rotor, which is quenched in non-viscous solvents but lights up in viscous media. To further characterize molecular rotor properties, we performed lifetime measurements (Figure 3C). The fluorescence lifetime of Th-R increased with viscosity. This is a typical behavior of molecular rotors which are quenched because of rotation in the excited state. However, one should note that the dependence was not linear, with rather slow growth of the lifetime at low viscosity range, and significantly steeper slope at higher viscosities.

**Figure 3.**
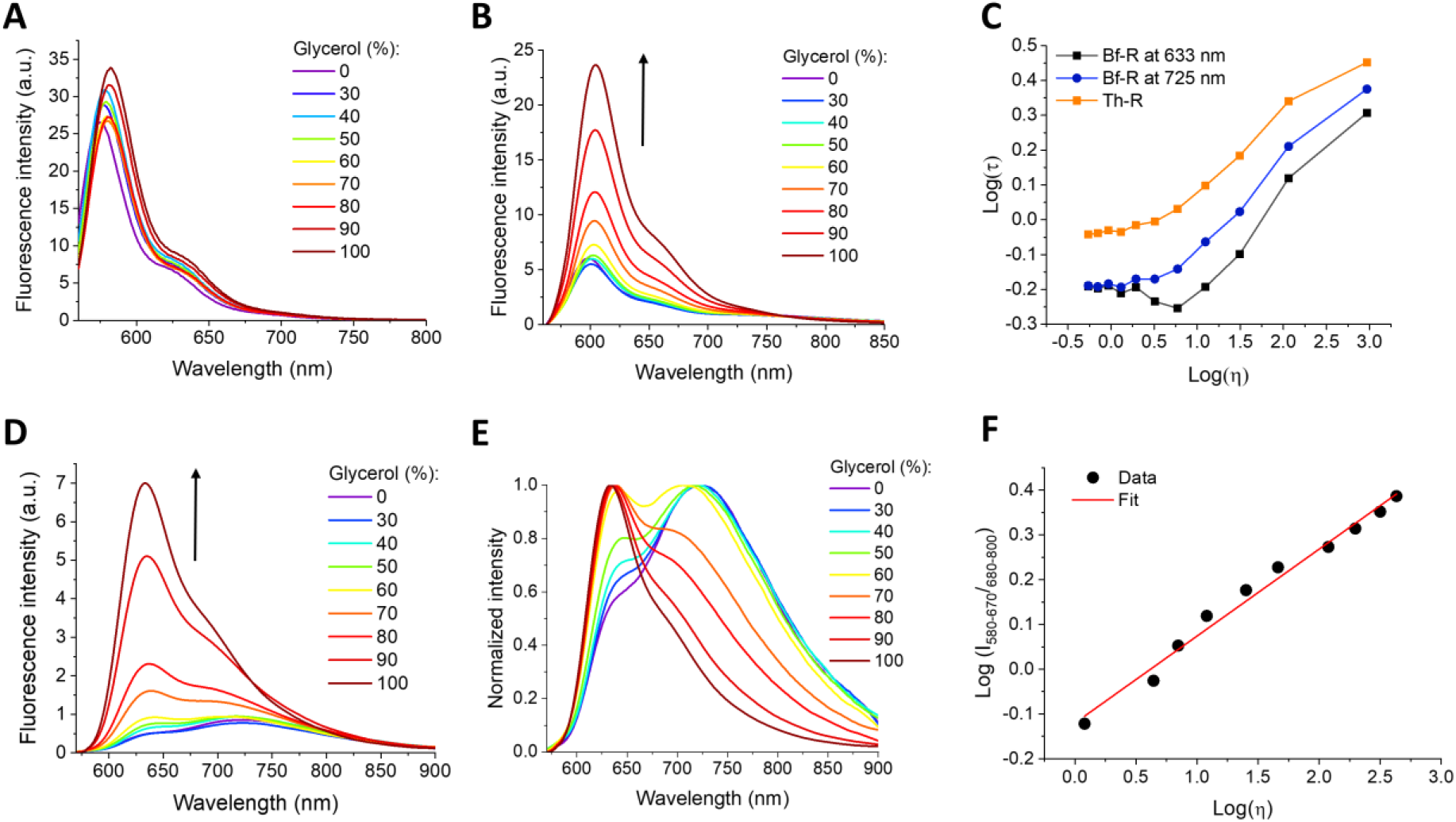
Fluorescence spectra of Ph-R (A), Th-R (B) and Bf-R (C) and normalized fluorescence Bf-R (D) in binary mixtures of MeOH and glycerol at varied volume fraction of glycerol (0-100%). (E) Logarithm of the intensity ratio vs logarithm of viscosity for the varied binary mixtures of MeOH and glycerol. (F) Correlation of fluorescence lifetime of Bf-R (black and wine curves) and Th-R with the viscosity. λ_ex_ = 550 nm.

Bf-R response to viscosity was associated with both intensity and spectral changes. Thus, increase in the viscosity resulted in the gradual increase in the intensity of the short-wavelength band, while the long-wavelength intensity almost did not change or slightly decreased (Figure 3D). The normalized spectra clearly showed dramatic changes in the relative intensities of the two emission bands as a function of viscosity (Figure 3E). Importantly, the plot of the logarithm of the intensity ratio of the red and NIR emission bands showed a linear relationship with viscosity over the entire range of viscosities studied (Figure 3F). This result suggests that Bf-R can be used to measure a broad range of viscosity values using simple ratiometric measurements. To the best of our knowledge this is the first example of molecular rotor based on a single fluorophore that exhibit ratiometric response to viscosity. Previous reports on probes with two-color ratiometric response to viscosity were based on two fluorophores.^30-31^ The fluorescence lifetime measurements also revealed clear dependence of fluorescence lifetime on viscosity of both emission bands (Figure 3C). The dependence showed a very similar trend as that for Th-R, which relatively small changes at low viscosities and a steep nearly linear dependence at higher viscosity values. This observation shows that the ratiometric and the lifetime outputs of Th-R do not follow exactly the same response profile, where ratiometric output showing much broader operation range with a linear response to viscosity.

To understand better the observed viscosity dependence of both Th-R and Bf-R, we performed DFT and TD-DFT calculations (Figure 4). First, we calculated the optimal geometry of the three dyes in the ground (S0) and excited (S1) states. For Ph-R, both the S0 and S1 states were found to be twisted, with 9-aryl/fluorophore dihedral angles of ∼58° and ∼53°, respectively. Moreover, the energy of the S1 state strongly depended on the dihedral angle value (Figure 4B), where planar conjugation (angle = 0) corresponded to very high energy unfavorable form (+12.5 kcal/mol). This means that for Ph-R the free rotation is practically impossible in the S1 state explaining the poor dependence of its emission on viscosity. Th-R also showed a twisted configuration in both S0 and S1 states, with angle values of ∼54° and ∼44°, respectively. However, its profile of energy vs dihedral angle was very different from Ph-R (Figure 4B), where flat configuration presented a much lower energy (∼2.6 kcal/mol), which suggests that this dye can undergo rotation of its aryl group. The latter can explain the appearance of molecular rotor properties of Th-R in comparison to Ph-R. The most remarkable results were obtained for Bf-R: it showed planar configuration in the S1, but a twisted one in S0 state (Figure 4A,B). In fact, its profile of energy vs dihedral angle was opposite to that of Th-R: low dihedral angles were preferred over higher angles, with the maximal energy around 90 degrees (∼3.5 kcal/mol). Thus, similarly to Th-R, Bf-R can also freely rotate in the excited state, which explains the dependence of its emission intensity on viscosity. Moreover, the difference in the geometry between S0 and S1 states may explain the emergence of the new emission band. Indeed, after electronic excitation to S1, Bf-R would undergo planarization with further emission to the planar ground state according to the Franck-Condon principle. This means that the dye will emit from the energetically stabilized S1 to energetically destabilized S0 state and therefore, this dye should present an S1→S0 transition with much lower energy gap and thus strongly red shifted emission. Our TD-DFT calculations predict a red shift of ∼0.2 eV (∼1500 cm^−1^) of the emission relative to the S1→S0 transition of the twisted configuration (Figure 4D). This prediction matches well to the position of the new red-shifted band in the NIR region observed for Bf-R. The additional long wavelength band for rosamine derivative was previously observed in low temperature glasses for the derivative with electron donor 9-aryl group, which was interpreted as bichromophore behavior, involving this group in the electronic excitation. This work supports the idea that implication of benzofuryl group in the electronic excitation of the dye is behind the appearance of the new long-wavelegnth emission band.^37^ However, in contrast to previous report, our probe provides dual emission at room temperature and with ratiometric response to viscosity.

**Figure 4.**
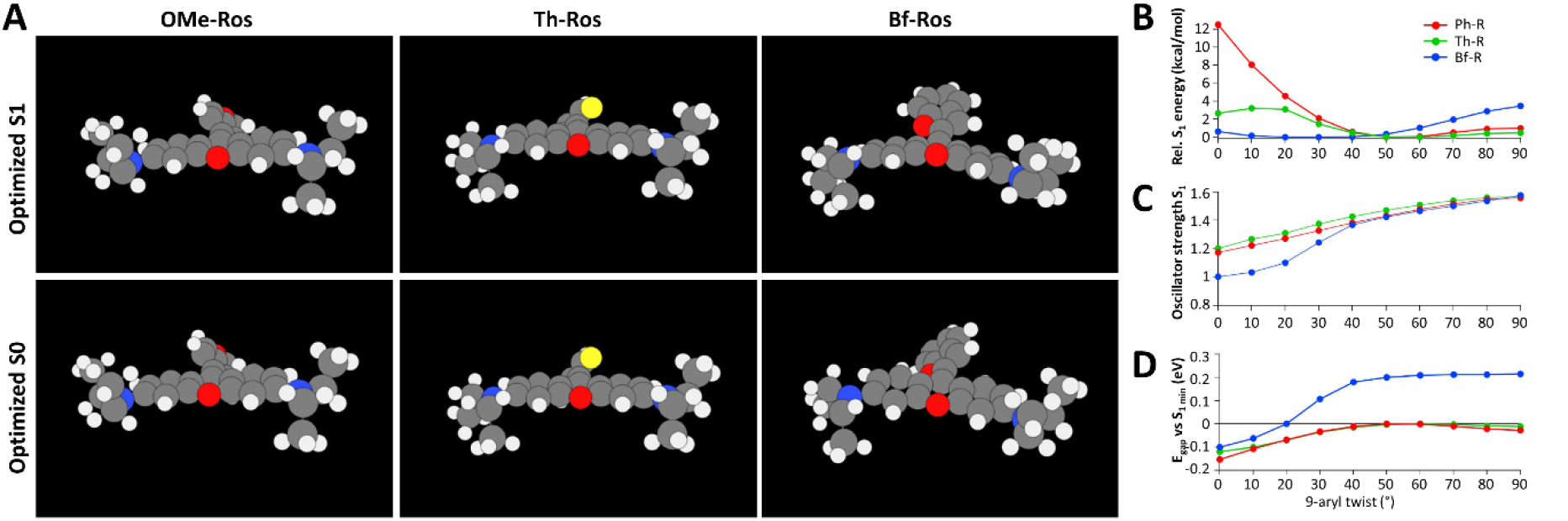
DFT and TD-DFT calculations for Ph-R, Th-R and Bf-R. Optimized geometries of the S0 (bottom panels) and S1 (top panels) states of the three dyes (A), together with the relaxed potential-energy profiles of the S1 state as a function of the meso-aryl dihedral angle (0° = coplanar, 90° = perpendicular, B). Ground-state geometries were optimized at the PBE0-D4/def2-SVP level and the S1 torsional profiles at CAM-B3LYP-D4/def2-SVP, with implicit solvation (CPCM). For Ph-R the S1 energy rises steeply toward planarity (+12.5 kcal/mol), so aryl rotation is essentially frozen; for Th-R the profile is shallow (∼2.6 kcal/mol), allowing rotation; for Bf-R the profile is inverted, with the S1 minimum near the planar geometry and twisting to 90° costing ∼3.5 kcal/mol. (C) Oscillator strength of the S1 state along the same coordinate: the S1 transition remains bright for all three dyes (f ≈ 1.0–1.6). (D) Change in the vertical S1→S0 energy gap (i.e. emission energy) relative to its value at the S1 minimum of each dye, as a function of the twist angle: for Ph-R and Th-R the emission energy varies only weakly (∼0.05 eV), whereas for Bf-R it changes strongly and monotonically being red-shifted near planarity and blue-shifted by ∼0.2 eV upon twisting.

However, this situation should be favorable mainly in non-viscous solvents, where the molecule can undergo relaxation to the planar S1 state. On the other hand, in the viscous media, the dye will probably not have time to relax, thus favoring emission of the twisted state (with an angle close to that of its S0 configuration). This state should have very similar characteristics to that for Th-R and its intensity should also grow with viscosity, in line with our experimental data. As a result, an increase in viscosity should favor emission from the twisted, blue-shifted configuration, whereas a non-viscous medium should favor emission from the planar configuration, which explains the increase in the red/NIR ratio with increase in the viscosity.

### Cellular experiments

Then, we studied the localization of Th-R and Bf-R after addition to the live cells (Figure 5). In U87 cells, both probes showed characteristic fibrillary structures, which co-localized with a mitochondria-labelling reference dye, Mito tracker Deep Red (Pearson’s coefficient is 0.96 and 0.97 for Th-R and Bf-R, respectively). In HEK and KB cell lines, both probes also showed very good colocalization with the Mito tracker Deep Red (Figure S1 and S2), suggesting that the new rhodamine derivatives are robust markers of mitochondria. The observed selective accumulation in the mitochondria was expected because their close analogues rhodamine derivative, like rhodamine 123 or tetramethylrhodamine methyl ester, are commonly used trackers of mitochondria,^39-40^ due to their net positive charge delocalized within the fluorophore. Our dyes present the same features, which make them capable to easily penetrate the cells and accumulate in the inner part of mitochondria membrane driven by the strong potential inside of the inner membrane.

**Figure 5.**
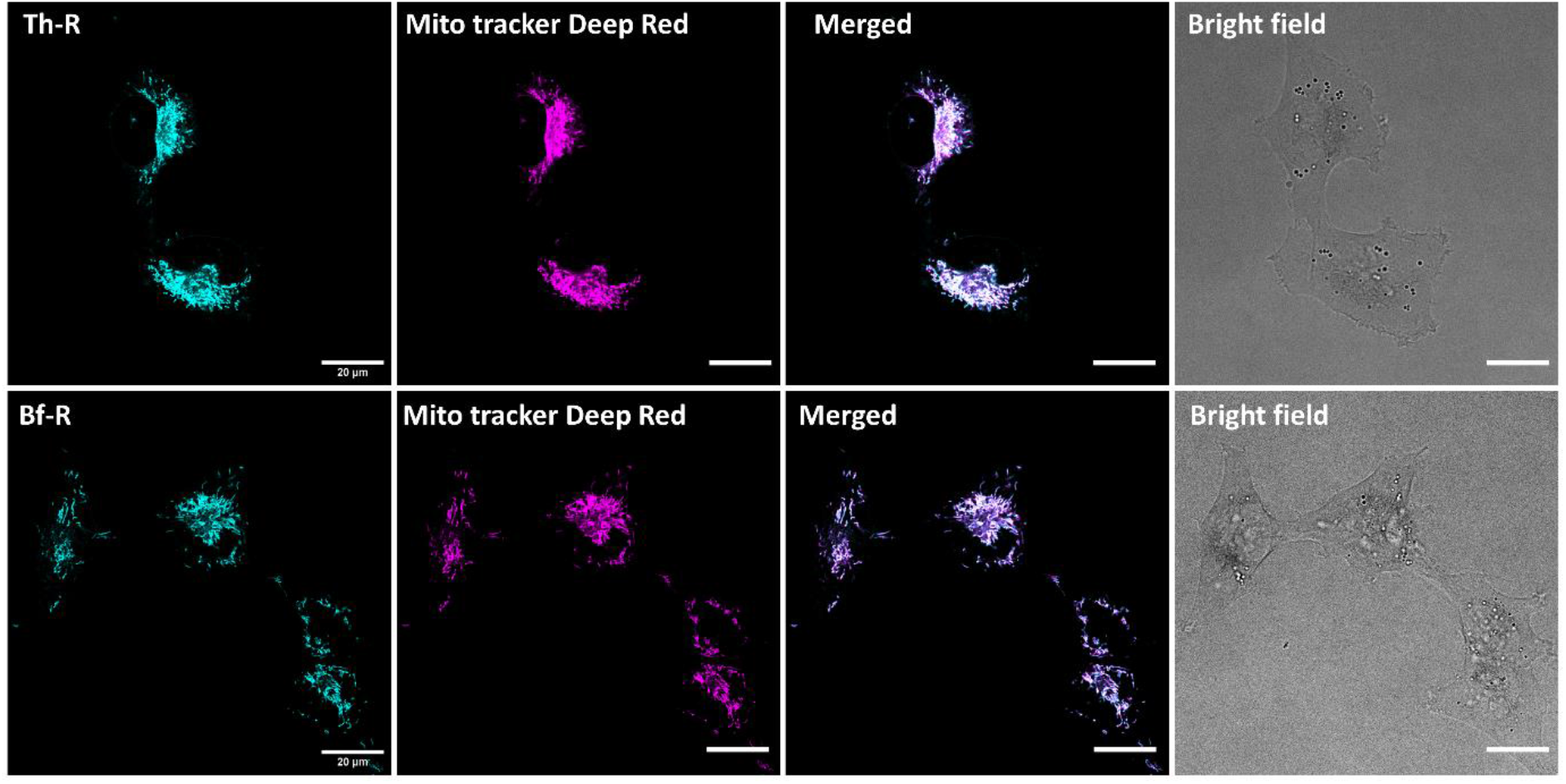
Fluorescence confocal imaging of U87 cells stained with Th-R or Bf-R (20 nM) together with mitochondria-labelling reference dye, Mito Tracker Deep Red (50 nM). Merged images and bright field images are also shown. Scale bars: 20 µm.

To understand better the nature of the local environment of Bf-R, we performed ratiometric imaging of the U87 cells stained with this dye. To this end, the emission was split into two channels: 580-670 and 680-800 nm, corresponding to the two emission bands of the probe. Then, the ratio of these two channels was built, which according to our studies showed correlation with environment viscosity. On the images, this ratio was represented as a pseudo-color of 16-color scale (Figure 6A). In parallel, we performed calibration of our probe under the microscope by recording images of solution of Bf-R in model mixtures of varied viscosity using the same microscope settings (Figure 6B). The individual calibrating solutions appeared homogeneous in terms of intensity and color. With increase in the viscosity, the pseudo-color of these solutions varied gradually from blue to red, which corresponded to the increase in the ratio values form 0.73 for 40% glycerol in methanol to 2.23 in pure glycerol (Figure 6C). On the other hand, the ratiometric image of the U87 cells showed a green-yellow pseudo-color of mitochondria, which corresponded to the ratio values of 1.40±0.05 (Figure 6C). The observed pseudo-color (and corresponding ratio value) is close to that of a calibrating solutions with 70-80% glycerol, corresponding to viscosity range between 250 and 460 cP.^41^ In HEK293T and KB cells, similar values of the intensity ratio were observed: 1.47±0.05 and 1.56±0.05, respectively (Figure S3). These results clearly suggest that our probe is located in a highly viscous medium, which is probably the inner membrane of mitochondria for all three cell lines. To verify this, we recorded the fluorescence spectra of Bf-R in the presence of large unilamellar vesicles (LUVs) composed of a phospholipid DOPC, which is a model of biological membranes. Importantly, in the presence of LUVs, Bf-R displayed ∼100-fold increase in fluorescence compared to the buffer (Figure S4), indicating that the probe can partition into lipid membrane. Therefore, its emission should reflect the viscosity in its membrane surrounding. At higher temperature (60 °C), the probe showed lower overall intensity and lower contribution of the short-wavelength band, as expected for less viscous membranes (Figure S4). The comparison of the obtained two-band spectrum at 20°C and the corresponding intensity ratio with those of model glycerol-methanol solutions (Figure 3E and S4) revealed that the local viscosity of the membrane corresponded to 70% glycerol solution (∼254 cP).^41^ This high value of viscosity in the model membrane matched remarkably well with that measured in mitochondria using the same probe. This result confirms our hypothesis that Bf-R senses viscosity in the inner membrane of mitochondria. The observed microviscosity values also matched well with the previous reported values the inner membrane of mitochondria, in the range 200-700 cP for NIH 3T3 cells^28^ and 275 cP in HeLa cells,^26^ estimated using other molecular rotor dyes.

**Figure 6.**
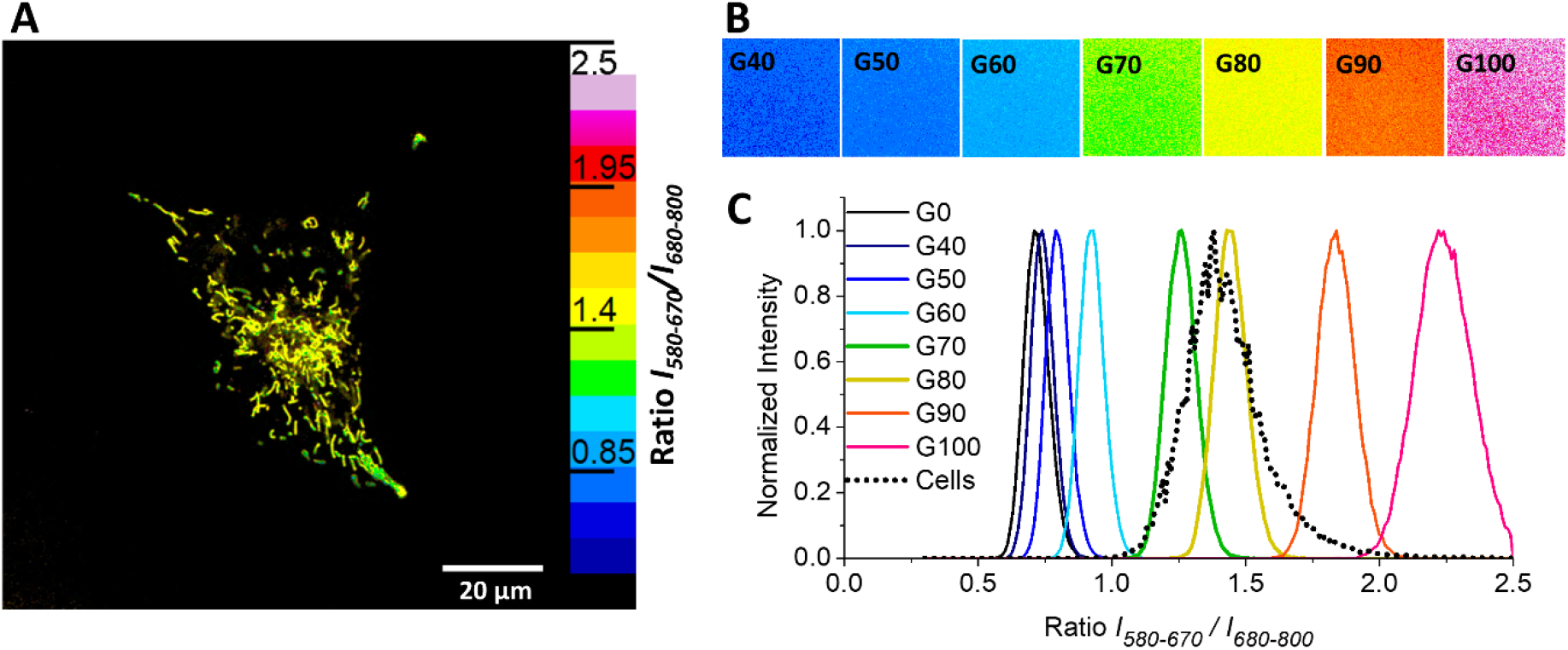
(A) Confocal fluorescence ratiometric image of U87 live cells stained with Bf-R (20 nM) and (B) calibration of the probe in the glycerol-methanol mixtures using the sample microscope settings. The same pseudo-color scale was used for (A) and (B). Scale bars: 20 µm. (C) Distribution of the intensity ratio values for methanol-glycerol mixtures and cells.

The capacity of the probe to target specifically mitochondria and probe its local viscosity prompted us to test a number of biological applications, in particular, the impact of the external stress on the local properties of mitochondria.

We tested the effect of oxidative stress produced by hydrogen peroxide (H_2_O_2_), which is known to profoundly impact mitochondria.^42^ According to our ratiometric images in U87 cells, the mitochondria stained with Bf-R changed the pseudo-color from green-yellow to yellow-orange, indicating the increase in the intensity ratio (Figure 7). The image quantification confirmed that the increase in the ratio, i.e. mitochondrial local viscosity was significant according to both single-cell analysis (Figure 6C) and the distribution of the ratio values (Figure S5). Very similar effects were also observed for HEK293T and KB cells (Figure S5, S6 and S7), indicating that the increase in the viscosity of the inner mitochondrial membrane after the oxidative stress produced by H_2_O_2_ is a generic phenomenon. The obtained result is in line with literature data that show systematic increase in microviscosity of the inner mitochondrial membrane in response to oxidative stress induced by H_2_O_2_^43^ or photodynamic therapy treatment.^44^

**Figure 7.**
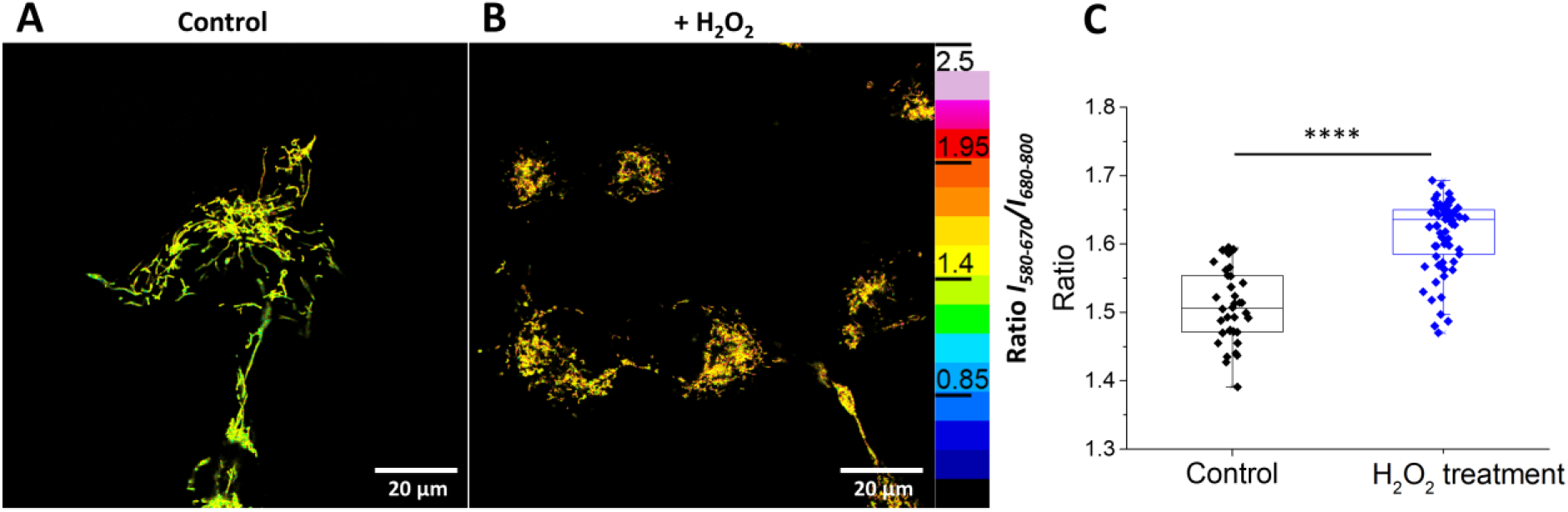
Fluorescence ratiometric images of U87 live cells without (A) and after (B) hydrogen peroxide treatment (2 mM H_2_O_2_ for 1h at 37 °C) with following Bf-R (20 nM) staining. Scale bars: 20 µm. (C) Single-cell quantification of the intensity ratio for these two conditions. ****p < 0.0001. Number of analyzed cells per condition n > 30.

## Conclusions

Fluorescent molecular rotors have become powerful tools for sensing local viscosity in biological systems and their applications spans from sensing lipid membrane organization and DNA packing to monitoring protein aggregation. However, the current molecular probes respond to viscosity by changing their fluorescence intensity and fluorescence lifetime, where the former is not convenient for quantification, while the latter requires specialized equipment. Ratiometric molecular rotors that change their emission color in response to viscosity would be an attractive alternative, but at this moment, their examples are limited to complex systems built from two chromophores. Here, we introduce a new design concept of ratiometric molecular rotor based on rhodamine dye. The latter is one of the most popular dye scaffolds, but it has never been used for designing ratiometric molecular rotors. Here, to develop this probe we explored the rotation of 9-aryl group of xanthene. To this end, three rhodamine derivatives were synthesized, where 2-carboxyphenyl group was replaced with 4-methoxyphenyl (Ph-R), 2-thienyl (Th-R) and 2-benzofuryl (Bf-R). We found that, in contrast to 4-methoxyphenyl derivative, 2-thienyl one exhibited strong dependence of its fluorescence intensity and lifetime on viscosity of solvent. Theoretical calculations confirmed that in the five-membered ring 2-thienyl can freely rotate in the excited state of the dye, which is not the case of 4-methoxyphenyl. Thus, 2-thienyl analogue appears as a classical molecular rotor with intensity and lifetime-based response to viscosity. On the other hand, benzofuryl derivative exhibits dual emission with the additional emission band in near-infrared region. This dye exhibits a fluorescence ratiometric response to viscosity, where the ratio of the short-wavelength to the long-wavelength bands increases with viscosity. Theoretical calculations suggest that the benzofuryl derivative can also freely rotate in the excited, similarly to thienyl. The major difference is that it prefers planar conformation in the excited S1 state, which could emit to less energetically favorable S0 state, which produces a new lower energy transition, explaining appearance of the NIR emitting band. Higher viscosities could block rotation of the 2-benzofuryl group, thus favoring short-wavelength emission of the dye from non-planar conformation, close to that of its ground states. Cellular studies by fluorescence microscopy revealed that both Th-R and Bf-R efficiently target mitochondria, which is in line with the behavior of cationic rhodamine dyes derivatives. Importantly, 9-benzofuryl derivative enables quantitative ratiometric imaging of local viscosity in mitochondria membrane, which revealed mitochondria heterogeneity with viscosity values ranging from 250 to 460 cP for U87 cells. This value corresponded well to local viscosity measured for Bf-R in model lipid membranes, which suggests that the probe senses viscosity of inner mitochondrial membrane. Oxidative stress produced by hydrogen peroxide treatment induced significant rise in the local microviscosity in addition to morphological changes, such as partial fragmentation of mitochondria. This result suggests that the probe can sense local chances in biophysical properties of mitochondria in response to external stress conditions. In this work, we propose a concept that convert bright rhodamine dyes into a molecules rotor with two-band ratiometric response to viscosity, which opens the route to simple quantitative sensing and imaging of viscosity in biological systems.

## Supporting information

Supplemental information for publication.

## Acknowledgments

This work was supported by Agence Nationale de la Recherche (ANR) NanoLipoVirus ANR-24-CE92-0026. We acknowledge PIQ platform of France-Bioimaging (FBI) network for providing access to fluorescence imaging facilities and PACSI platform − for NMR and mass spectrometry.

