## Supplemental information for publication. for "Rhodamine-derived ratiometric fluorescent molecular rotor for mitochondrial viscosity sensing"

### Materials and Methods

**General.** All reagents and solvents were purchased from suppliers (Sigma-Aldrich, TCI, Fisher Scientific) of analytical grade and used as received. Milli-Q-water (Millipore) was used in all experiments. NMR spectra were recorded at 20 °C on a Bruker Avance III 400 MHz and 500 MHz spectrometer for $^1$H and $^{13}$C, respectively. Absorption and fluorescence spectra were recorded on a UV-2700i UV-VIS spectrophotometer (Shimadzu) and FS5 spectrofluorometer (Edinburg Instruments), correspondingly. All experiments were performed at room temperature. Generally, fluorescence quantum yields (QY) determination was done using rhodamine 101 (R101) as a standard ($QY_{MeOH} = 1$). The 35 mm dishes with a glass coverslip bottom were purchased from ibidi® and LabTek™ Chamber Slide (borosilicate glass, eight-wells) was purchased from ThermoFischer Scientific for microscopy.

#### Synthesis and characterization

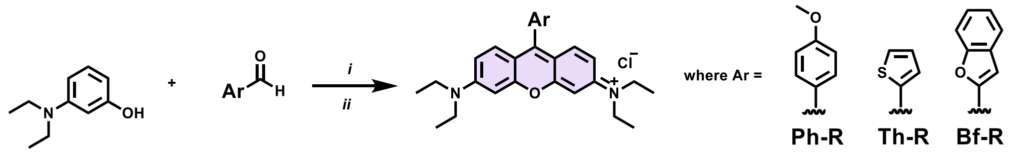


**Scheme S1**. Synthesis of rhodamine-based molecular rotors: i) DCM:MeOH (10:1), TFA, rt, overnight; ii) chloranil, DCM:MeOH (1:1), 4Å molecular sieves, rt, overnight.

**General procedure for synthesis of rosamine derivatives.** 3-(Diethylamino)phenol (2 equiv) and appropriate benzaldehydes or heterocyclic aldehydes (2 equiv) were placed in a round bottom flask and were dissolved in mixture of DCM:MeOH (10:1), 50 mL and 5 mL respectively. After 500 µL of TFA was added. The reaction mixture was stirred overnight at room temperature (RT) under inert atmosphere. Reaction was followed by TLC. When reaction was completed, solvents were removed under the vacuum and the crude product was used for the next step. The crude oil was dissolved in DCM:MeOH (1:1), 60 mL and 60 mL, respectively, followed by addition to reaction mixture of chloranil (1.1 equiv) and 5 volume % of 4Å molecular sieves. The reaction mixture was stirring overnight at RT. Upon completion, the solvents were evaporated under reduced pressure. Purification was performed by column chromatography using DCM:MeOH (98:2 to 9:1) as a mobile phase.

**N-(6-(diethylamino)-9-(4-methoxyphenyl)-3H-xanthen-3-ylidene)-N-ethylethanaminium chloride (Ph-R)**. Yield 30 %. $^1$H NMR (400 MHz, $CDCl_3$) δ (ppm): 7.46 (d, 2H), 7.36 (d, 2H), 7.15 (d, 2H), 6.91-6.89 (dd, 2H), 6.86 (d, 2H), 3.94 (s, 3H), 3.66 (m, 8H), 1.35 (t, 12H). HRMS-ESI (*m/z*): $[M]^+$ calculated for $C_{28}H_{33}N_2O_2$ 429.255, found 429.2546.

**N-(6-(diethylamino)-9-(thiophen-2-yl)-3H-xanthen-3-ylidene)-N-ethylethanaminium chloride (Th-R)**. Yield 25 %. $^1$H NMR (400 MHz, $CD_3OD$) δ (ppm): 7.98 (d, 1H), 7.72 (d, 2H), 7.48 (d, 1H), 7.42 (m, 1H), 7.13-7.10 (dd, 2H), 6.95 (d, 2H), 6.85 (d, 1H), 6.33-6.28 (m, 3H), 3.72-3.66 (q, 8H), 1.34 (t, 12H). HRMS-ESI (m/z): [M]+ calculated for C25H29N2OS 405.1993, found 405.1989.

**N-(9-(benzofuran-2-yl)-6-(diethylamino)-3H-xanthen-3-ylidene)-N-ethylethanaminium chloride (Bf-R).** Yield 20 %. $^1$H NMR (400 MHz, DMSO-$d_6$) δ (ppm): 7.97-7.9 (m, 3H), 7.84-7.79 (m, 2H), 7.59 (m, 1H), 7.48 (m, 2H), 7.23 (d, 2H), 6.98 (s, 2H), 3.69 (m, 8H), 1.25 (t, 12H). HRMS-ESI (m/z): $[M]^+$ calculated for $C_{29}H_{31}N_2O_2$ 439.2381, found 439.2376.

**Fluorescence Lifetime Measurements.** Lifetime measurements of Th-R and Bf-R in MeOH/glycerol mixtures were performed using Time-Correlated Single Photon Counting (TCSPC) technique. Excitation pulses at 550 nm were generated by NKT Photonics SuperK Extreme laser with-repetition rate 10 MHz. The fluorescence decays were collected at 570 nm for Th-R, 633nm and 725nm for Bf-R, respectively, using a polarizer set at magic angle and a 16 mm band-pass monochromator (Jobin Yvon). TCSPC system is equipped with an ultrafast microchannel plate photomultiplier tube detector (Hamamatsu R3809U-51) and electronics board (Becker & Hickl SPC-130) and has instrument response time about 60-65 picoseconds. Triggering signal for the TCSPC board was generated by sending a small fraction of the laser beam onto a fast (400 MHz bandwidth) Si photodiode (Thorlabs Inc.). Fluorescence signal was dispersed in Acton Research SPC-500 monochromator after passing through a pump blocking, long wavelength-pass, autofluorescence-free, interference filter (Omega Filters, ALP series). The monochromator is equipped with a CCD camera (Roper Scientific PIXIS-400) allowing for monitoring of the time-averaged fluorescence spectrum. A stock solution of Th-R and Bf-R with concentration 100 μM were prepared in DMSO. For each 0-100% glycerol/MeOH mixture the corresponding dye was added to achieve an optical density around 0.1 in absorption spectra.

**Formulation of Liposomes.** Large unilamellar vesicles (LUVs) of various compositions have been made using previously described method[1]. Briefly, aliquots of stock solutions DOPC (5 mM), cholesterol (10 mM), and SM (10 mM) in $CHCl_3$ have been placed into 10 mL round-bottom flask and solvent was evaporated under vacuum. After all, 10 mL of PB (20 mM, pH = 7.4) was added in order to get 1 mM lipids concentration. Then a suspension of multilamellar vesicles was filtered through 0.2 μm (7 passages) and 0.1 μm (10 passages) using a Lipex Biomembranes extruder (Vancouver, Canada). The final filtered formulations of LUVs were characterized in terms of size and polydispersity index with Zetasizer Nano ZSP (Malvern Panalytical, U.K.). The molar ratio of lipid composition was 1:0.9 in a case of DOPC/Chol and SM/Chol, respectively.

**Cell Culture and Treatment Conditions.** HEK 293T cells (add provider) were grown in Dublecco's Modified Eagle Medium (DMEM, Gibco) supplemented with 10% fetal bovine serum FBS (Gibco), 1% L-glutamine (Sigma-Aldrich), 1% penicillin-streptomycin (Gibco) at 37 °C in a humidified 5% CO2 atmosphere. U87 and KB cells were grown in Minimum Essential Medium (MEM, Gibco) and supplemented with 10% FBS (Lonza), 1% L-glutamine (Sigma-Aldrich), 1% penicillin-streptomycin (Gibco), 1% non-essential amino acids (Gibco), % phenol-red (Gibco) at 37 °C in a humidified 5% CO2 atmosphere. For live-cell imaging studies, cells were seeded on 35 mm glass-bottom dishes at a density $5\times10^4$ cells/well 24 h before microscopy measurements. The medium was removed and remained attached cells were washed three times with Hank's Balanced Salt Solution (HBSS, Gibco). Next, staining solution of dyes in HBSS was added to the cells.

*Colocalization Experiments.* Cells were co-incubated with Th-R or Bf-R and commercial Mito TrackerTM Deep Red (Invitrogen) for 30 min at 37 °C and then confocal imaging was performed without any washing steps. Working concentration of dyes was 20 nM and 50 nM for Mito tracker.

*Oxidative Stress.* Cells were incubated with 2 mM H2O2 for 1 h at 37 °C to induce oxidative stress. After cells were washed one time with HBSS followed of incubation with 20 nM Bf-R for 30 min at 37 °C. Untreated cells were used as a control.

*Effect of Nystatin.* Cells were passed onto surface of μ-dishes at density 1.5□105 cells/well. Next day nystatin treatment was applied to cells with working concentration of nystatin 10 μM followed of 1 h incubation at 37 °C. After all, the remaining attached cells were washed one time with HBSS and Bf-R was added keeping 20 nM concentration and incubation for 30 min at 37 °C. Correspondently, cells without nystatin addition were used as a control.

**Confocal Microscopy.** Live-cell imaging was performed on a Leica laser scanning confocal microscope (TSC SP8) at room temperature. Images were obtained using 552 nm and 638 nm lasers for rhodamine dyes and for Mito-tracker, respectively. The fluorescence was detected at two spectral ranges 580-670 nm and 680-800 nm for Bf-R, in the range 560-610 for Th-R, and 648-685 nm for Mito-tracker.

### Supporting figures

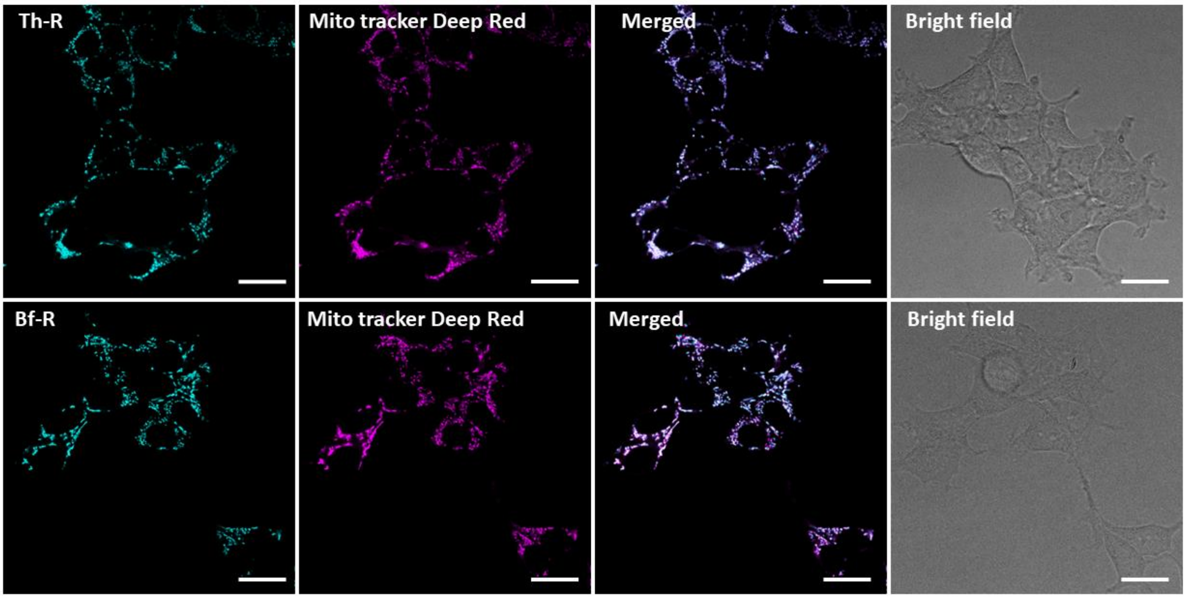


**Figure S1.** Fluorescence confocal imaging of live HEK cells stained with Th-R or Bf-R (20 nM) together with mitochondria-labelling reference dye, Mito tracker Deep Red (50 nM). Merged images and bright field images are also shown. Scale bars: 20 µm.

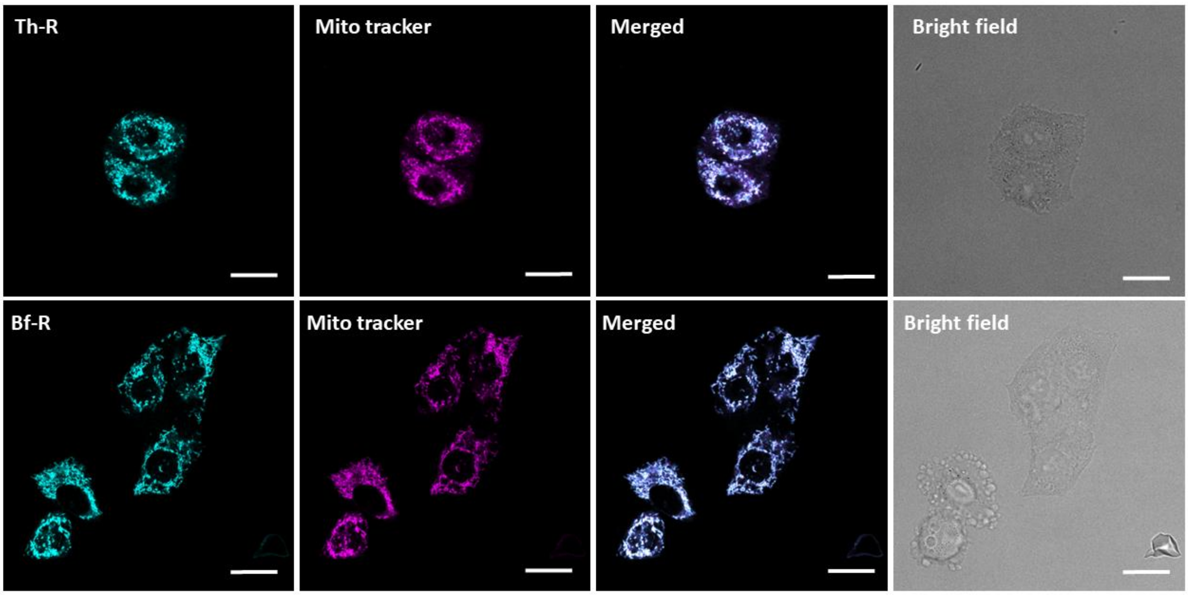


**Figure S2.** Fluorescence confocal imaging of live KB cells stained with Th-R or Bf-R (20 nM) together with mitochondria-labelling reference dye, Mito tracker Deep Red (50 nM). Merged images and bright field images are also shown. Scale bars: 20 µm.

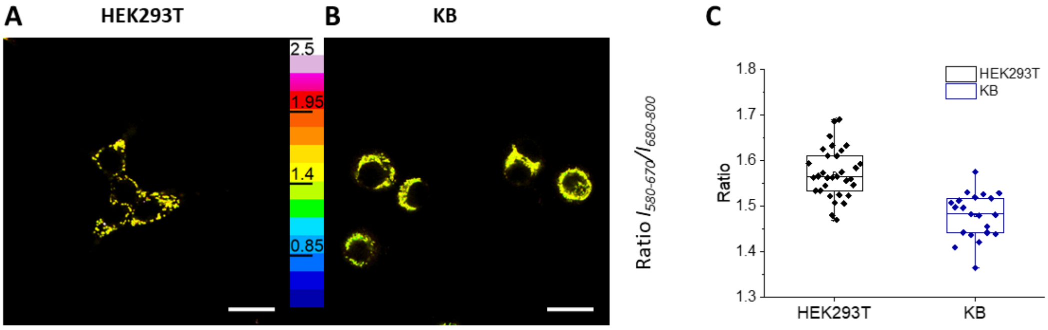


**Figure S3.** Fluorescence ratiometric imaging live HEK293T (A) and KB (B) cells after staining with Bf-R (20 nM). Scale bars: 20 µm. (C) Single-cell quantification of the intensity ratio for HEK293T (black dots) and KB cells (blue dots). Number of analyzed cells per condition n > 20.

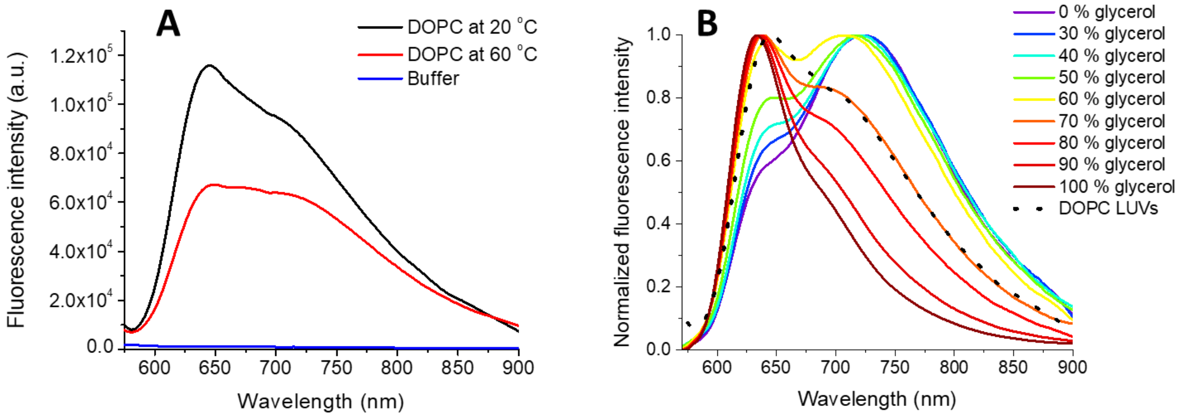


**Figure S4**. (A) Fluorescence spectra of Bf-R in phostaphe buffer (20 mM, pH 7.4) and with DOPC vesicles (0.2 mM lipid) at two temperatures and (B) comparison of the spectrum at 20 °C vs those recorded for glycerol-methanol mixtures.

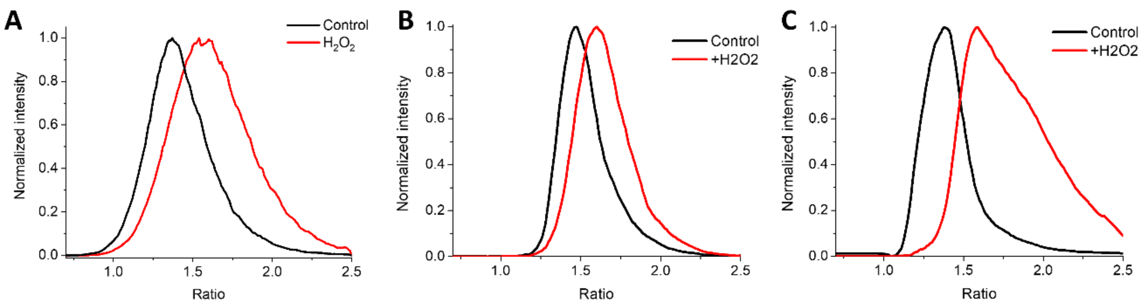


**Figure S5**. Distribution of the intenisty ratio obtained according to the analysis of the ratiometric images of U97 (Figure 6), HEK (Figure S6) and KB (Figure S7) cells.

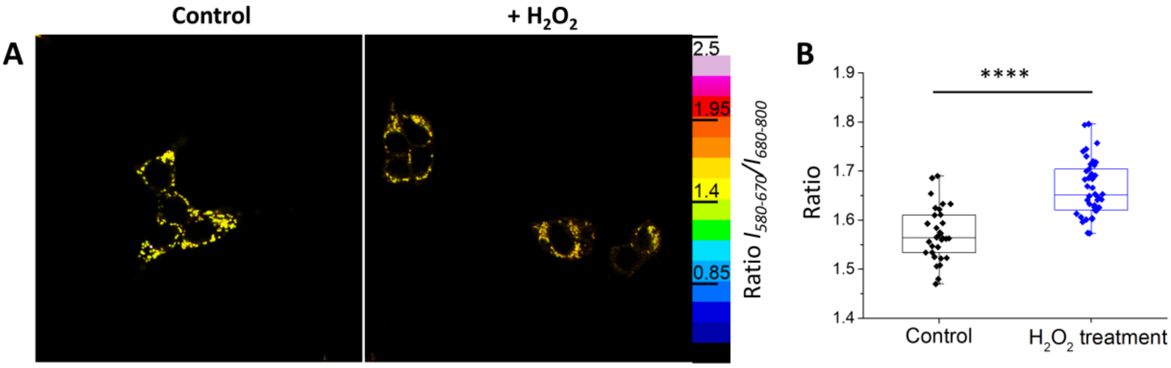


**Figure S6**. Fluorescence ratiometric images of live HEK293T cells without (A) and after hydrogen peroxide treatment (2 mM $H_2O_2$ for 1h at 37 °C) followed staining with Bf-R (20 nM). Scale bars: 20 µm. (B) Single-cell quantification of the intensity ratio for for these two conditions. ****p < 0.0001. Number of analyzed cells per condition n > 25.

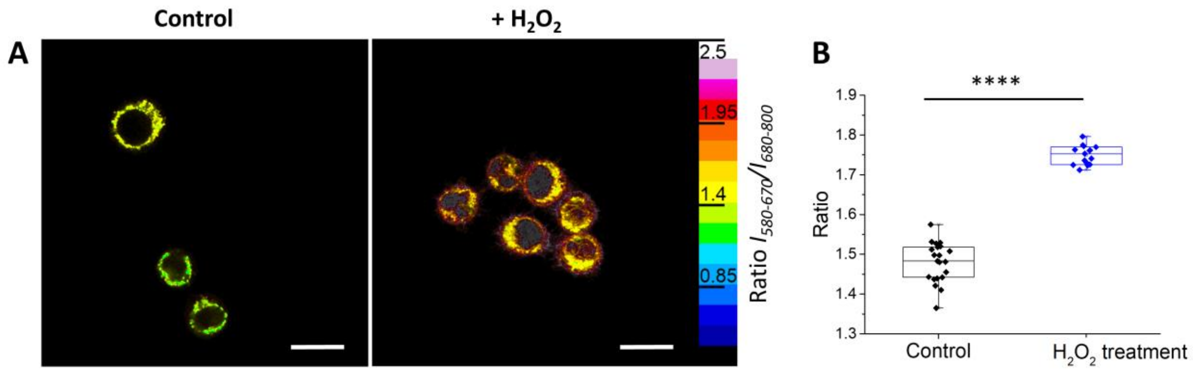


**Figure S7**. Fluorescence ratiometric images of live KB cells (A) without and after hydrogen peroxide treatment (2 mM $H_2O_2$ for 1h at 37 °C) followed staining with Bf-R (20 nM). Scale bars: 20 µm. (B) Single-cell quantification of the intensity ratio for these two conditions. ****$p < 0.0001$. Number of analyzed cells per condition $n > 12$.
